# Interplay between cell proliferation and recruitment controls the duration of growth and final size of the Drosophila wing

**DOI:** 10.1101/2021.05.14.444212

**Authors:** Elizabeth Diaz-Torres, Luis Manuel Muñoz-Nava, Marcos Nahmad

**Affiliations:** Department of Physiology, Biophysics, and Neurosciences, Center for Research and Advanced Studies of the National Polytechnic Institute (Cinvestav-IPN), Avenida Instituto Politecnico Nacional 2508, Colonia San Pedro Zacatenco, Mexico City, 07360, MEXICO

**Keywords:** growth control, robustness, cell recruitment, cell proliferation, mathematical modeling, Drosophila wing disc

## Abstract

How organs robustly attain a final size despite perturbations in cell growth and proliferation rates is a fundamental question in developmental biology. Since organ growth is an exponential process driven mainly by cell proliferation, even small variations in cell proliferation rates, when integrated over a relatively long time, will lead to large differences in size, unless intrinsic control mechanisms compensate for these variations. Here we use a mathematical model to consider the hypothesis that in the developing wing of Drosophila, cell recruitment, a process in which undifferentiated neighboring cells are incorporated into the wing primordium, determines the time in which growth is arrested in this system. Under this assumption, our model shows that perturbations in proliferation rates of wing-committed cells are compensated by an inversely proportional duration of growth. This mechanism ensures that the final size of the wing is robust in a range of cell proliferation rates. Furthermore, we predict that growth control is lost when fluctuations in cell proliferation affects both wing-committed and recruitable cells. Our model suggests that cell recruitment may act as a temporal controller of growth to buffer fluctuations in cell proliferation rates, offering a solution to a long-standing problem in the field.

**Graphical abstract:** 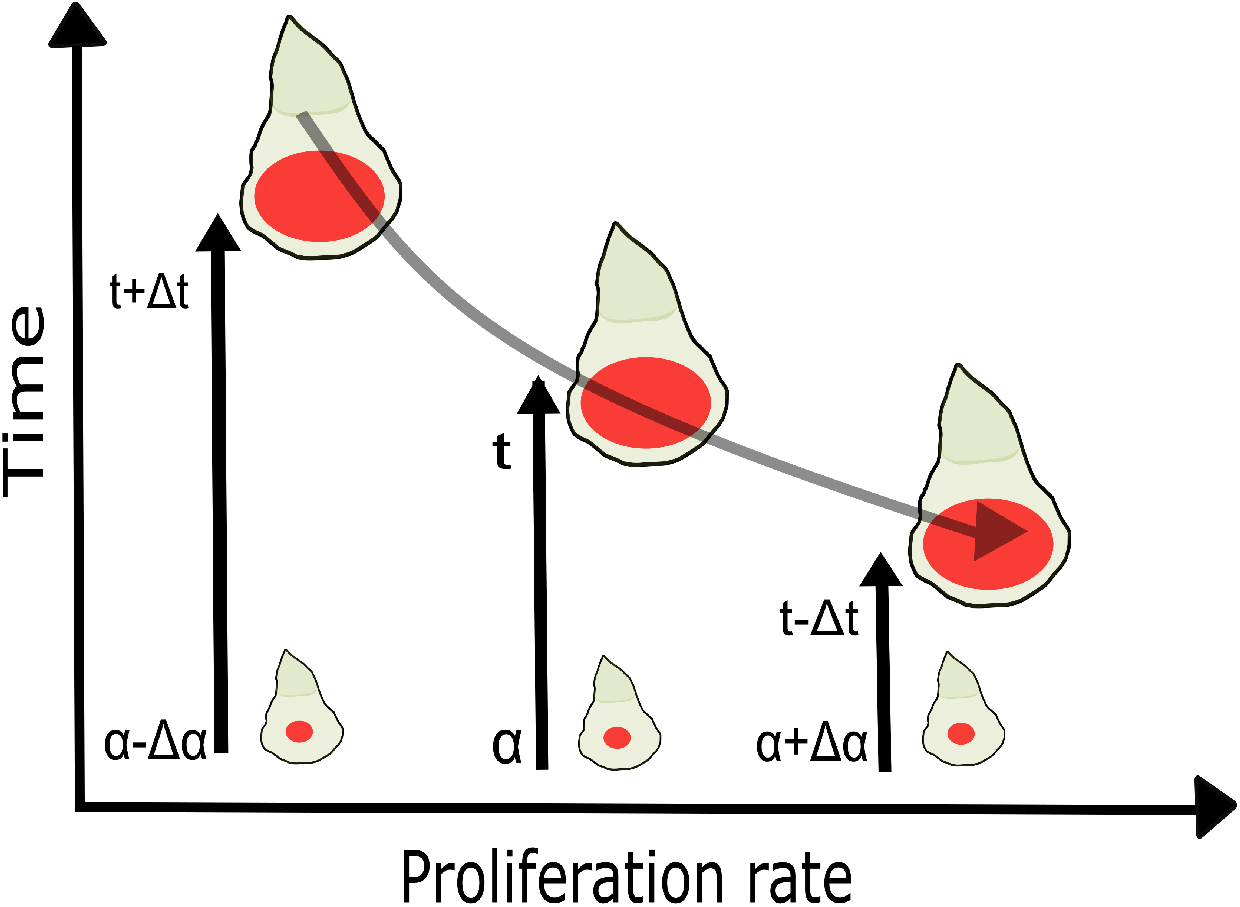

## 1. Introduction

During animal development, both extrinsic and intrinsic cues contribute to the final size of organs [1]. Extrinsic cues such as nutrition, hormones, and temperature drive changes in overall body size, allowing organisms to adapt to different environmental conditions and energetic resources, while maintaining functional organs [2]. While size adaptation to genetic fluctuations and changing environments is an important driver of phenotypic evolution, organisms also require a robust control of organ size in order to ensure their proper function. Understanding how body plans are designed to permit variability under certain perturbations while robustly reaching a constant organ size under other perturbations is a fundamental problem in developmental biology [3].

The Drosophila wing disc is a useful model to investigate the mechanisms underlying developmental patterning and organ growth [4,5,6,7,8]. Evidence suggests that chemical gradients (morphogens), nutrition, hormones, and mechanical interactions contribute to extrinsic and intrinsic growth, but how all these signals are integrated to result in a wing of a specific size and shape is little understood [9,10,11,12,13]. While extrinsic factors determine *the* ultimate size of the wing, transplantation experiments of wing discs into the abdomen of an adult fly or experiments in which pupation is halted and the duration of larval development is increased, show that discs are able to stop growing when a particular size is reached [14,15,16]. Moreover, the fact that wing discs stop growing at the ‘right’ size even when cell proliferation, cell sizes, or cell numbers are perturbed suggest that there are intrinsic mechanisms that robustly control the final size of the disc [17,18]. Several hypotheses have been proposed to explain how wing stop growing at the right size, including the morphogen slope model [19] and the mechanical feedback model [20, 21,22]. However, experimental evidence in support of these models remains inconclusive.

The wing disc of *Drosophila* grows exponentially during three larval instars, mainly through cell proliferation and, to a lesser extent, through cell recruitment, a process by which undifferentiated neighboring cells are incorporated into the wing tissue by the induction of the wing selector gene, *vestigial* (*vg*) [23,24,25]. This process occurs only during the third larval instar and contributes to about 20% of adult wing size (including proliferation of newly recruited cells) [26]. However, it is unknown whether there is a crosstalk between cell proliferation and recruitment to coordinate growth and achieve the proper wing size.

Because cells in the wing disc divide every 6-10 hours over a period of four days [27], even small perturbations in cell proliferation rates would result in large differences in final size [28,29]. Here we investigate whether cell recruitment works as a fine-tuning mechanism to buffer fluctuations in cell proliferation rates and control adult wing size. Using a mathematical model, we explore the dynamics of populations of cells within the wing pouch and find parameter sets that agree with a time-course analysis of wing-committed *vs*. *recruitable* cells. We show that growth by cell proliferation accelerates the rates of cell recruitment by increasing the number of *recruiter* cells. In particular, we show that recruitable cells extinguish at a time that is inversely proportional to variations in cell proliferation rates. Taken together, we propose that cell recruitment acts as a temporal controller that allows wings to attain a specific final size despite perturbations in cell proliferation rates.

## 2. Material and Methods

### 2.1 Image visualization, processing and data analysis

All the images were processed and analyzed using ImageJ/Fiji software (https://imagej.net/), and the matplotlib (https://matplotlib.org/), pandas (https://pandas.pydata.org/), NumPy (http://www.numpy.org/), SciPy (https://www.scipy.org/) Python packages.

To quantify the W and R cells within the wing pouch, we used the immunofluorescence images stained with Vg, Wingless (Wg), and DAPI in y, w (considered wild type in this study) discs used in a previous publication [26]. For initial conditions W_0_ and R_0_, we took early third instar images and counted manually (twice by two different observers). To compute R_0_, we subtracted the number of nuclei (marked by DAPI) within the inner ring of Wg to the number of nuclei stained with Vg (W_0_; data not shown). For the quantification in Fig. 3, the wing pouch of every wing disc was determined in the DAPI channel using the ellipse selection tool in ImageJ, taking as reference for the minor axis the folds that limit this region in the ventral and dorsal edges. As these folds have a curve like shape, we manually adjusted the ellipse to fit these curve lines and indirectly obtained the major axis. To remove the background noise of Vg, we selected a rectangle in a region of the wing discs where this protein is not expected to be present (outside the pouch) and made an average of the intensities within this area to have a unique threshold value to determine whether a pixel’s Vg intensity is background noise. After obtaining this threshold we binarized all the pixels within the wing pouch in having or not having Vg protein. To compute W and R cells, we first remove the pixels not corresponding to the nuclei of Vg-expressing cells. Using the Particle Analysis tool of ImageJ to define the Vg positive pixels within a nucleus, we eliminated the pixels marked as Vg positive but isolated from other Vg+ ones (we reasoned that these pixels were marked due to non-specific binding of the antibody). The size parameter of this tool was set to 50 (i.e., we quantified that in average 50 pixels formed the cell’s nucleus), the circularity parameter between 0 and 1 to admit different nuclei shapes and the holes between nuclei were also stained as Vg positive to avoid counting them as R pixels. Finally, all the resulting pixels within the wing pouch positive to Vg correspond to the W cells and the negative ones to the R cells.

### 2.2 Optimization procedure

For the optimization problem in Fig. 4C, we considered couples of parameters (*ρ, k*) within the black area of parameter space in Fig.4C, that defines those parameter sets that correspond to a *t_f_* within the range of 24 and 36 hours, which is the time in which the recruitment process takes place [26]. For each parameter set (*ρ, k*), we perturbed α (only α_W_ in Fig. 2D,E; and both α_W_ and α_R_ in Fig. 2F,G) within the range of [0.75α_0_, 1.25α_0_], where α_0_ is the wild-type value reported in Table 1. For each perturbation, we then solve equations (3.1–3.2) and computed the final time (*t_f_*) defined by the time in which *R* → 0 and the recruitment rate ε, computed as described in Fig. 2B. We then compared *t_f_* with the final time (*t_fctl_*) that would leave the final size W_f_ invariant (see Fig. 4C, right) and computed the difference between these values for each perturbed *α*. We defined the optimal parameter set as such that minimizes this difference (Fig. 4D, F). We noted that this control works for the range of perturbations [0.75α_W0_, 1.25α_W0_]. When this range is significantly increased, there are significant variations in *t_f_* — *t_fctl_* (Sup. Fig. 3). The worst fit (*i.e*., the set that maximizes this difference) is shown in Sup. Fig. 4.

**Table 1.**
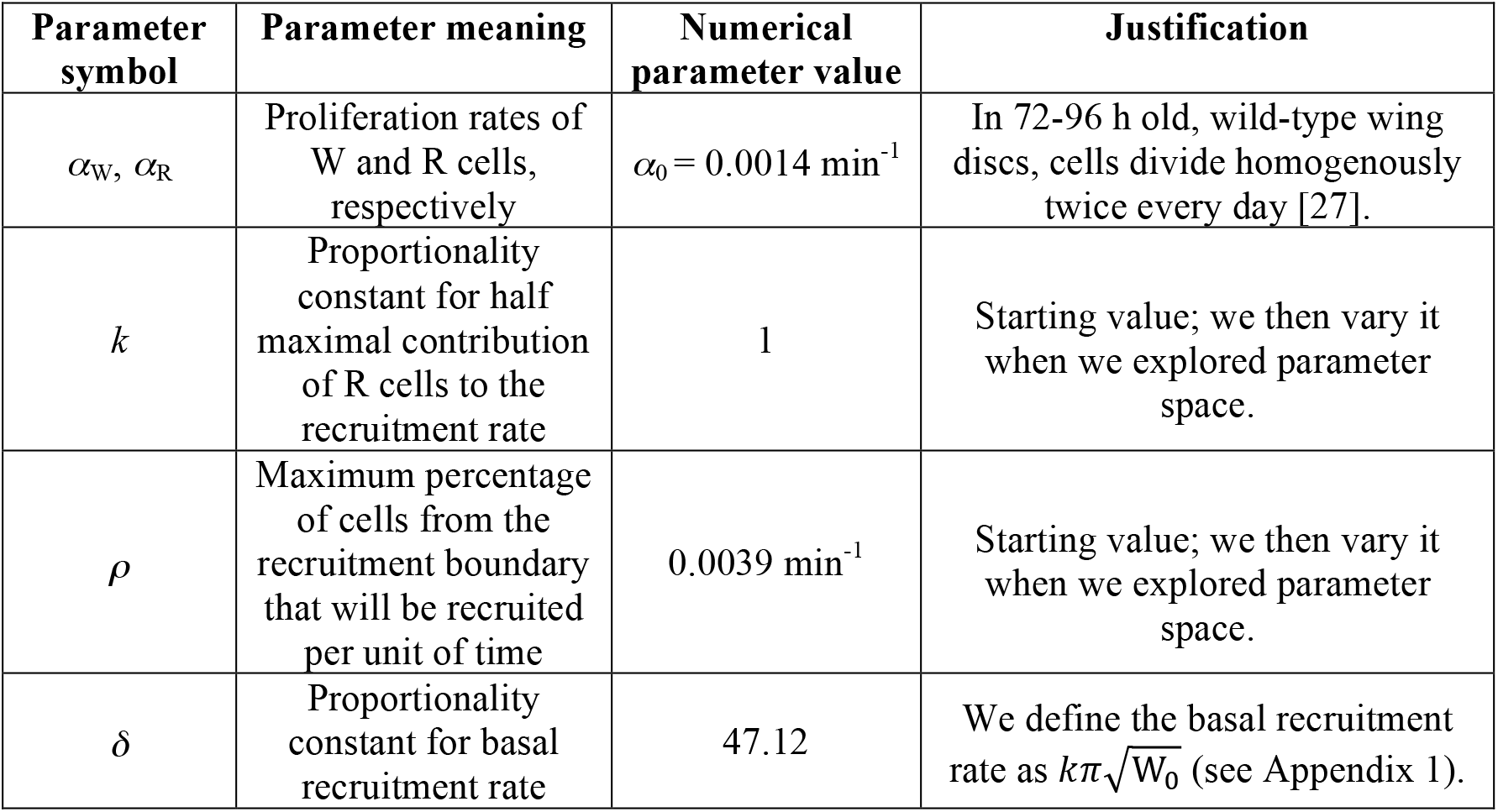

| Parameter symbol | Parameter meaning | Numerical parameter value | Justification |
| --- | --- | --- | --- |
| $\alpha_W, \alpha_R$ | Proliferation rates of W and R cells, respectively | $\alpha_0 = 0.0014 \text{ min}^{-1}$ | In 72-96 h old, wild-type wing discs, cells divide homogenously twice every day [27]. |
| $k$ | Proportionality constant for half maximal contribution of R cells to the recruitment rate | 1 | Starting value; we then vary it when we explored parameter space. |
| $\rho$ | Maximum percentage of cells from the recruitment boundary that will be recruited per unit of time | $0.0039 \text{ min}^{-1}$ | Starting value; we then vary it when we explored parameter space. |
| $\delta$ | Proportionality constant for basal recruitment rate | 47.12 | We define the basal recruitment rate as $k\pi\sqrt{W_0}$ (see Appendix 1). |

## 3. Theory

### 3.1 Construction of a mathematical model that incorporates cell proliferation and recruitment in the developing wing of Drosophila

We built a dynamical model of the populations of wing-committed, Vg-expressing cells (referred as W cells) *vs*. recruitable, pre-wing cells (referred as R cells), defined as the cells located within the wing pouch that do not express Vg at the moment, but are primed to express it through the recruitment signal [25] (Fig. 1A). We begin our simulations at the beginning of the third instar (about 80 h after egg laying). At this time, it is possible to manually count the numbers of W and R cells (W_0_ and R_0_, respectively; see Material and Methods). The model takes into account cell proliferation in each cell population (at rates *α*_W_ and *α*_R_) and cell recruitment (function F(W,R); Fig. 1B), which we assume is a radial and contact-dependent process that positively contributes to W, but at the same rate reduces R. F(W,R) depends on the geometry of the system and the number of cells located at the recruitment front (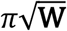; Appendix 1). We assume that cell death is negligible [30] and therefore we ignore it in the model. Under these assumptions, the dynamics of W and R are given by the following equations (Fig. 1B):

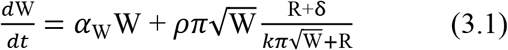

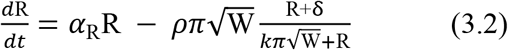

where *ρ* (rate units), δ and *k* (dimensionless) are constants of proportionality (see Table 1). Since cell proliferation is nearly homogeneous throughout the disc (except at the DV border) [27], we assume that *α*_W_ = *α*_R_, unless when proliferation is perturbed in a particular population of cells (see Results). Unperturbed proliferation rate will be referred below as *α*_0_ for both W and R populations. Parameter values are either extracted from previous experimental studies or explored numerically (see Table 1 and Results).

**Figure 1.**
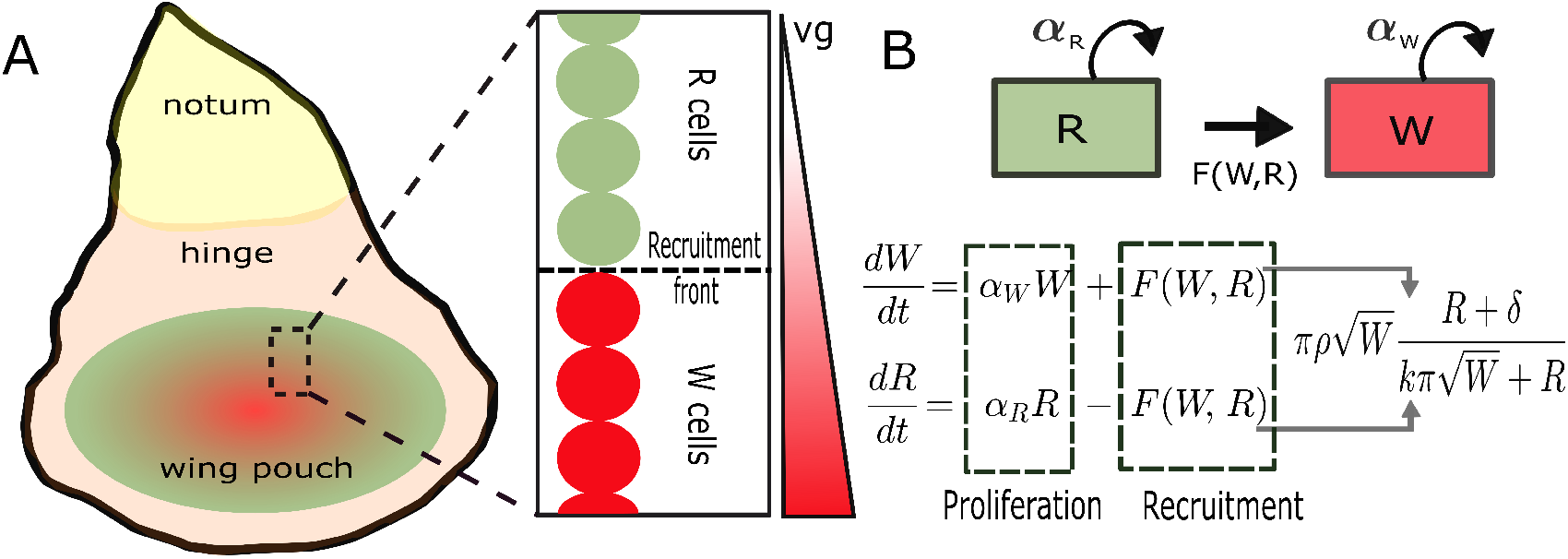
Scheme of the Drosophila wing disc and mathematical model of wing and recruitable cells. (A) Cartoon of a Drosophila wing disc, subdivided in three regions (notum, hinge, and wing pouch) that correspond to different structures in the adult. The wing blade derives from the wing pouch domain and may be subdivided in wing-committed (W) and pre-wing recruitable (R) cells. W cells are defined by the expression of Vg (red); as the disc grows, the Vg domain expands at the expense of reducing the R domain (green). (B) Dynamic model for the populations of W and R cells. Note that in addition to cell proliferation of W and R cells, the W population grows at the expense of the R population through a recruitment rate function F(W,R); see Appendix 1.

### 3.2 Other assumptions of the mathematical model

The model described by equations (3.1–3.2) is a valid representation of the dynamics of W and R cells under the following considerations. Notice that 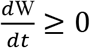 for all *t, i.e*., W is an increasing function of *t*. However, we attempt to investigate a final-state value of W at the time in which the population of R cells vanishes (referred below as final time, *t_f_*). This assumption is justified by the experimental observation that cells stop proliferating further after all the cells in the pouch are forced to express Vg, resulting in smaller wings [31]. Another assumption is that the introduction of the parameter *δ* assumes that there is a basal recruitment rate when the number of R cells becomes too small. If we do not make this assumption, the recruitment rate artificially decrease when R is too small (even when there are recruiter cells sending a recruitment signal). But of course, introducing *δ* causes R to become negative for *t*>t_f_. Thus, we assume that the model capture the population dynamics of W and R cells only for 0>*t*>t_f_.

## 4. Results

### 4.1 Growth by cell proliferation and cell recruitment of the population of wing-committed (W) cells is approximately exponential

We used our mathematical model to explore the relative contributions of cell proliferation and cell recruitment to the wing at the final time *t_f_*. Although the system of equations is nonlinear, it is possible to show analytically that W is bounded by an exponential function of time (Fig. 2A; see Appendix 2). Thus, we wonder what would the exponential contribution of cell recruitment be relative to the cell proliferation rate *α*_W_. In a semi-log plot, the slope of the best-fit line corresponds to approximate exponential growth rate of the W population (Fig. 2B):

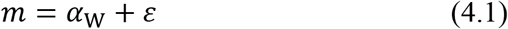

where *ε* is approximately the exponential grow rate due to cell recruitment. Thus, the final size of the W population is approximately given by (Fig. 2C):

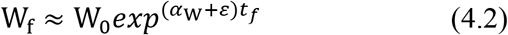

**Figure 2.**
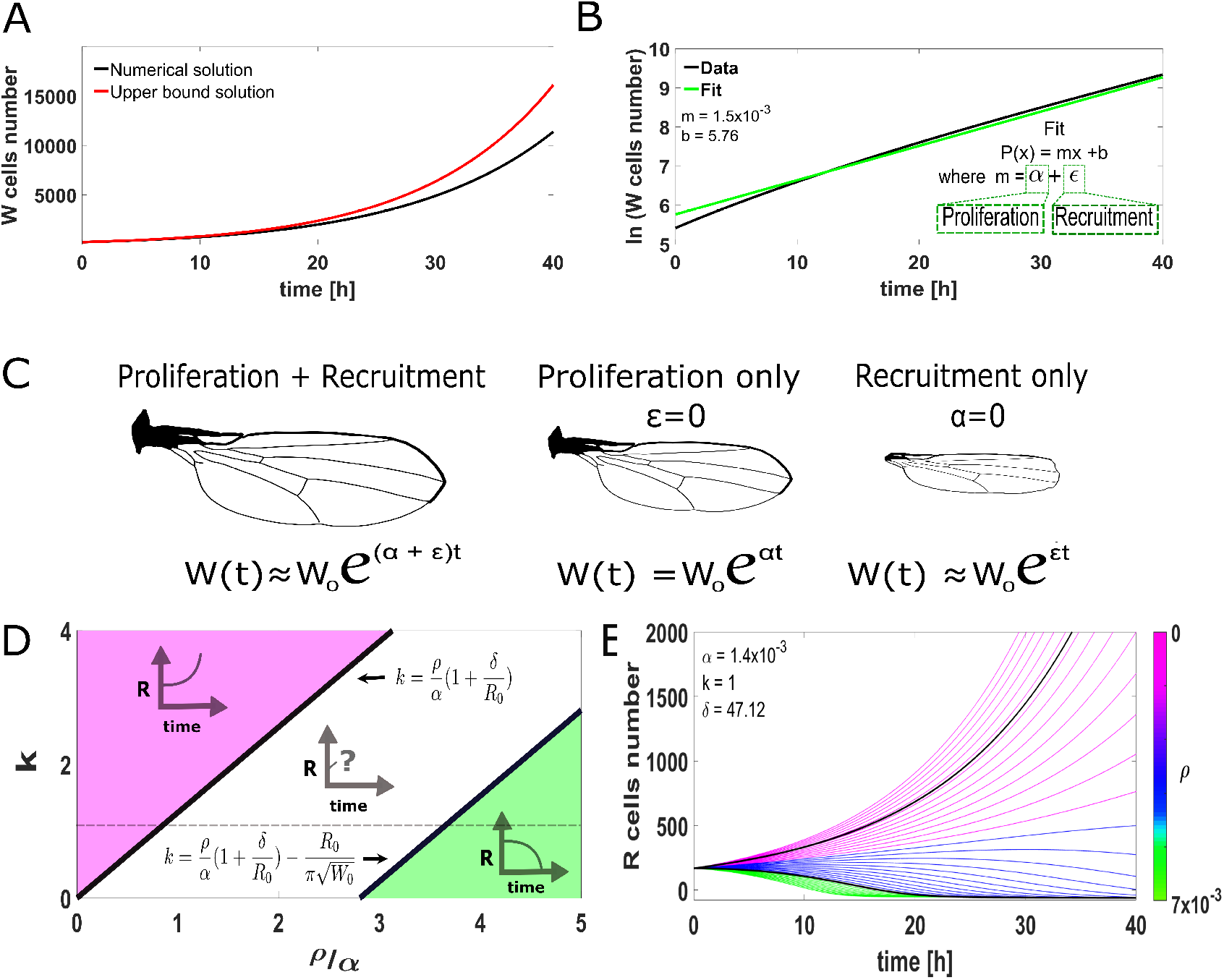
Dependence of W and R dynamics on system parameters. (A) Numerical solution of W from equations (3.1–3.2) using the parameters in Table 1 (black curve), compared with an analytical upper bound function (red curve); see Appendix 2. (B) Semi-log representation of the numerical solution in A (black curve) compared with the best fit solution of the linear function P(x)=mx+b. Note that because this is a semi-log plot, m represents the exponential rate of growth that has a contribution of proliferation (α_W_) and recruitment (ε). (C) Under the approximation in B, proliferation (*α*) and recruitment (*ε*) rates additively contribute to exponential wing growth. (D) The dynamics of R can be studied analytically in three different regions of parameter space; see Appendix 3. The region in pink corresponds to parameter values in which R is an increasing function of time (cell proliferation of R cells dominates over recruitment). Conversely, the region in green corresponds to parameter values in which R is a decreasing function of time (recruitment dominates over cell proliferation of R cells). The full dynamics of R in the white region cannot be defined analytically, but R is initially increasing. The dotted line represents a subset of parameters defined by *k*=1. (E) Numerical solutions for the population of R cells when *k, δ*, and *α* are fixed, and *ρ* is varied (color bar) along the dotted line in D. The dynamic behaviors found in D are reproduced numerically (black curves correspond to parameter values on the black lines in D). Note that within the white region, some solutions are increasing (as in the pink region), but some only increase transiently and then start to decrease (blue curves). The value of *ρ* reported in Table 1 corresponds to the middle value of line segment defined by the dotted line intersected by the two black lines in D.

A numerical estimate of ε (using the parameter values reported in Table 1) is 1.0×10^-4^, about 1/10 of the proliferation rate. This relative recruitment rate integrated over the expected duration of the recruitment process during third larval instar (36 h) is consistent with our previously-reported relative contribution of cell recruitment of about 20% [26].

### 3.2 Analytical and numerical exploration of parameter space in the model define the dynamics of the population of recruitable (R) cells

While W is always an increasing function of time, the dynamics of R is expected to depend on the parameters of the model. In particular, if cell proliferation of the R population dominates recruitment (*α*_R_>>*ρ*), we would expect R to be a continuously increasing function of time, a situation that does not recapitulate wild-type development as Vg eventually covers the whole wing pouch at the end of the third larval instar ([26]; see Fig. 3). Conversely however, if recruitment dominates cell proliferation in R cells then the R population will eventually vanish. We explored analytically the specific dependence of this dynamics on parameter values and initial conditions for the particular case in which *α* = *α*_R_ = *α*_W_, a case that appears to be a good approximation under wildtype conditions ([27]; see Appendix 3 for details). We found 3 stereotypical behaviors for the dynamics in parameter space (Fig. 2D). First, when 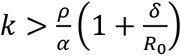 we find that R is an increasing function of time (pink region in Fig. 2D); in this case a final state is not reached and both W and R grow indefinitely, a condition that is not observed in the wild-type scenario. Second, when 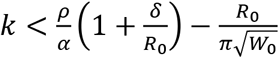 we found that R is a decreasing function of time (green region in Fig. 2D). Finally, when 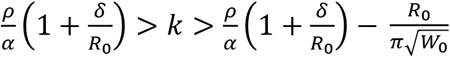, we were not able to demonstrate analytically the full dynamic behavior of R, but we show that 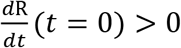, *i.e*., R is initially an increasing function of time (white region in Fig. 2D). We then explore numerically the dynamics in this region of parameter space by fixing *k*=1 and *α*=0.0014 and varying *ρ* (dotted line in Fig. 2D). We found that for some parameter values R is an increasing function of time for all *t*, while for others it increases for some time and then it begins to decrease (Fig. 2E, blue lines).

**Figure 3.**
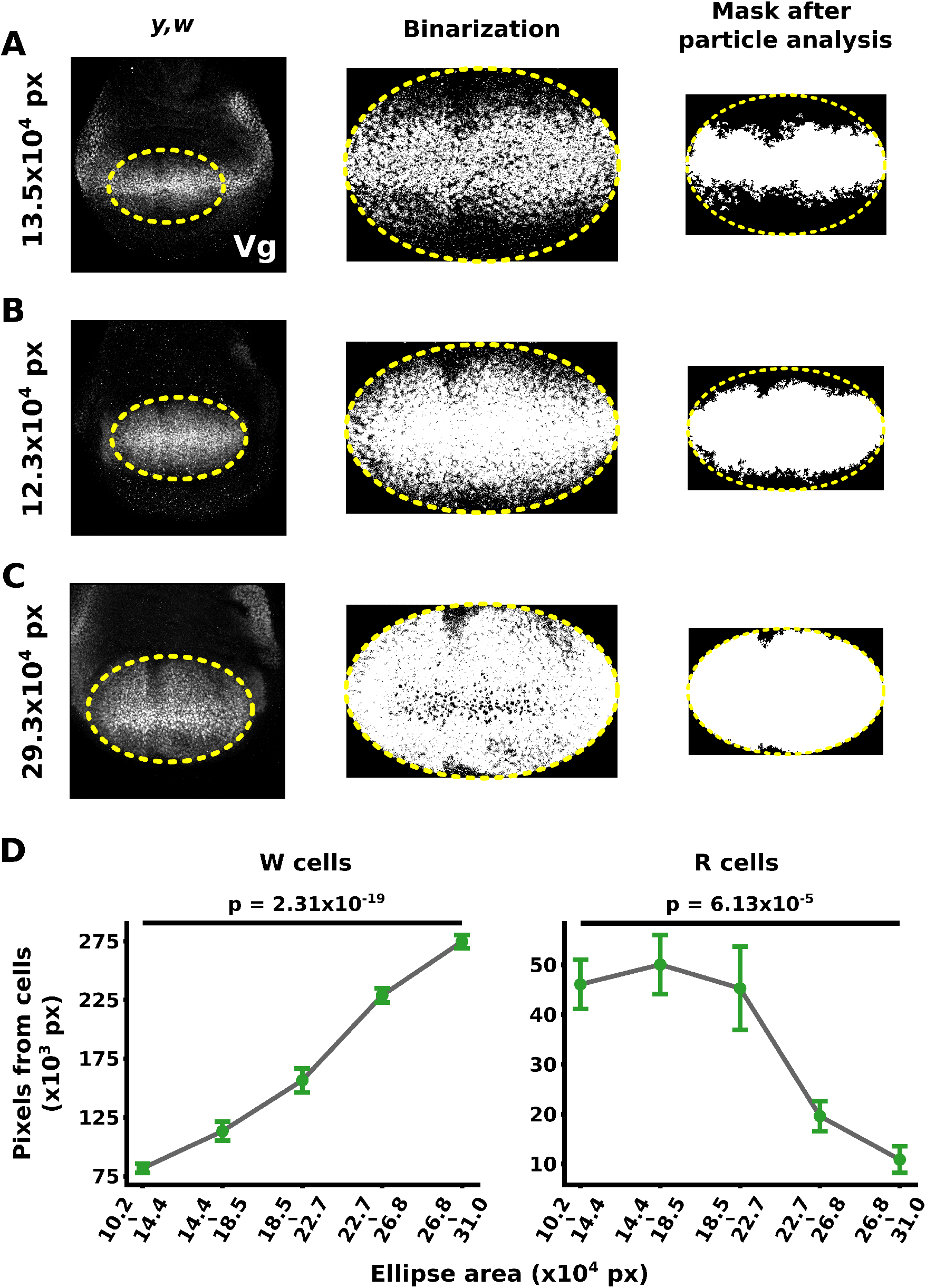
Quantification of W and R cells population in mid-to-late third instar wing discs. (A – C, left) Vg antibody staining in y,w (wild-type) discs of increasing sizes. The yellow dotted line delineates the contour of an ellipse that is chosen as the wing pouch area (see Material and Methods). (A – C, middle) Binarization of the pixels with or without Vg staining located within the wing pouch area (see Material and Methods). (A – C, right) Noise reduction of the Vg+ isolated pixels (error of the technique) using a particle analysis algorithm (see Materials and Methods). (D) Quantification of pixels representing W (left) and R (right) cells in discs of different sizes. Statistical test is an ANOVA one-way test.

### 4.3 Time-course analysis of Vg-expressing cells in the wing disc reveals that the population of recruitable (R) cells decreases by the end of the third instar

In order to validate the dynamics of W and R in our model with wild-type experimental data, we quantified pixels from cells in discs at different times during the third instar (Fig. 3A-C). While W cells are defined by Vg expression, unfortunately, there is no known molecular marker of R cells. Therefore, we used the absence of Vg expression in cells within the pouch to quantify R cells (see Material and Methods). As expected, we found that W increases exponentially and that R eventually decreases with time (Fig. 3D,E). In addition, our data suggests that the number of R cells appears to slightly increase or remain constant during the early stages of recruitment before it starts to decrease (Fig. 3E). This suggests that in wild-type development, the dynamics of R recapitulates the behavior of our mathematical model and identifies a specific region of parameter space where this behavior occurs (white region in Fig. 2D, blue curves in Fig. 2E). Interestingly, the fact that R does not decrease immediately appears to allow the recruitment process to continue throughout much of the third instar.

### 4.4 If growth is arrested when the population of recruitable cells is extinguished, perturbations in cell proliferation rates are compensated by the duration of growth

We then turned to explore the interplay between cell proliferation and recruitment with our mathematical model. First, we noticed that the recruitment function F(W,R) in Fig. 1B is an increasing function of time (Fig. 4A). We realized that this is due to the fact that as the population of W cells increases exponentially, the number of recruiter cells (which for a circular geometry is given by 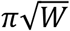) also increases exponentially, resulting in the acceleration of the recruitment process. This acceleration of recruitment not only happens as development progresses, but will be affected by perturbations of the proliferation rate *α*_w_. In fact, the amount of increase in F(W,R) responds proportionally to changes in *α*_0_, the wild-type value of *α*_w_ as reported in Table 1 (Fig. 4A,B). Therefore, our model suggests a natural way in which cell recruitment is fueled by cell proliferation rates. Because experimental evidence supports that growth is arrested when R→0 [31], we hypothesized that the coupling between cell proliferation and recruitment may work as a size control mechanism if upon a perturbation in cell proliferation rates, cell recruitment is accelerated or slowed down in a way that it terminates recruitable cells in a time that is inversely proportional to the growth rate perturbation (Fig. 4C). We asked if such control mechanism could work in the system for a set of parameters within those that match the dynamical behavior in Fig. 3E (black region in Fig. 4C). To test this, we performed an optimization procedure to search for parameter pairs (*k, ρ*) that minimize the error with the following optimal control function (see Fig. 4C, right; see Material and Methods):

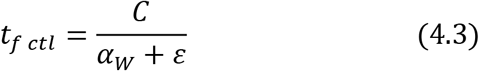

where *α*_w_ and *ε* correspond to the perturbed proliferation and recruitment rates, respectively, and *C* = (*α*_w0_ + *ε*_0_)*t*_*f*0_, is a constant that was chosen so the optimal control function coincides when at the unperturbed value of α_w_ (Fig. 4C, right). Remarkably, the optimal parameter values result in a function *t*_f_ that fits very well with the optimal control function defined by equation 4.3 (Fig. 4D). As a result, the final size of the W population, represented by W_f_, is mostly unaffected by small perturbations in *α*_w_ (Fig. 4E). Taken together, our model poses the hypothesis that cell recruitment could work as a controller of final size upon perturbations in cell proliferation rates by modulating the duration of growth in the opposite direction of the perturbation in cell proliferation.

**Figure 4.**
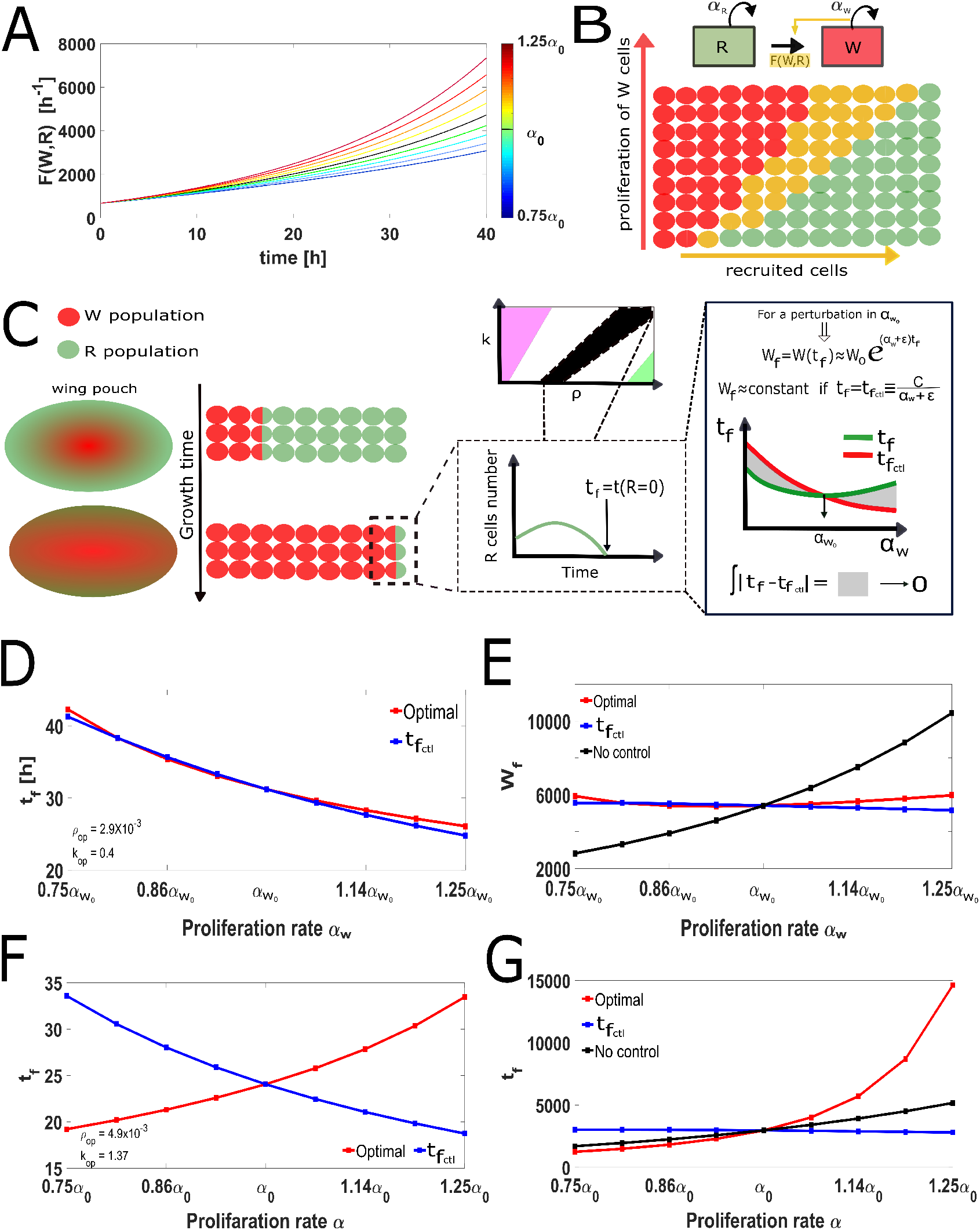
Coupling of cell proliferation and recruitment rates control the duration of growth and final size. (A) Numerical examination of F(W,R) with the parameters reported in Table 1, but when *α* is perturbed 25% above and below the wild-type value *α*_0_. Note that F(W,R) is always an increasing function of time, but the magnitude of the perturbation is proportional to the change in the recruitment rate. (B) Diagram illustrating the result in A. W cells are represented in red and R cells represented in green. The recruitment of new cells (yellow) increases as the proliferation rate of W cells increase. (C, left) Diagram illustrates that the propagation of the recruitment front is accelerated with developmental time. (C, center) There is a range of parameters in which R cells are extinguished within 24-36 h from the beginning of recruitment (black region). We define this extinction time as *t_f_*. (C, right) Optimization scheme that compares how *t_f_* compensates to perturbations in *α*_W_ in ideal conditions (red curve) *vs*. the compensation offered by the model (green curve). Assuming the exponential approximation of Fig. 2B, the *t_f_* that leaves the final size Wf invariant is such that is inversely proportional to the growth rate *α_W_+ε* (see Materials and Methods for details). (D) Plots of the optimal *t_f_* (red curve) and *t_fctl_* (blue curve; defined in panel C) *vs*. perturbations in *α*_W_. (E) Plots of the final number of W cells, W_f_, *vs*. perturbations in *α*_W_ corresponding to optimal *t_f_* (using the numerical solution of the model; red curve) and *t_fctl_* (using the equation for W_f_ in panel C; blue curve). The black curve corresponds to the numerical solution of the model when the optimal parameters are used, but *t_f_* is fixed to the wild-type final time *t*_*f*0_. (F) Same as D, except that now the optimization is attempted when both *α*_W_ and *α*_R_ (labeled as simply *α*) are perturbed simultaneously around the wild-type value of *α*_0_. Colors are defined as in D. Note that even in the optimal case, *t_f_* is an increasing function of perturbations in *α*_0_, (G) Same as E, except that now the optimization is attempted when both *α*_W_ and *α*_R_ (labeled as simply *α*) are perturbed simultaneously around the wild-type value of *α*_0_. Colors are defined as in D. Note that the red curve grows even faster than the black curve (no control).

### 4.5 Control is broken when cell proliferation in both W and R cells is perturbed

Experimental testing of our size control hypothesis may be challenging because impairing cell recruitment in R cells and, at the same time, introducing cell proliferation perturbations is not an easy task even in a powerful genetic model such as the Drosophila wing disc. In order to test our hypothesis, it would be useful to find a perturbation in which the growth control mechanism fails such that the model would predict very distinct phenotypes. In our perturbation simulations, we realized that recruitment works as a controller only when perturbations in cell proliferation affect W cells, but not R cells. Indeed, when perturbations affect both *α_w_* and *α_R_* simultaneously, we obtained the complete opposite behavior of *t_f_*, compared to the case in which only *α_w_* is affected (compare Fig. 4D and Fig. 4F). This is because increasing *α_R_* produce more R cells, so it would take longer to extinguish them by cell recruitment; and exactly the opposite happens when *α_R_* decreases. As a result of this, W_*f*_ is very sensitive to perturbations in proliferation rates (Fig. 4G). Thus, our model would predict very different behaviors when cell proliferation is perturbed only in W cells than when it affects both W and R cells. These contrasting phenotypes provide a relatively simple output to validate or reject our growth control model experimentally.

## 5. Discussion

Cell recruitment has been recently recognized as a patterning-driven growth mechanism in a plethora of systems [32], but whether there is a specific role in developmental growth is unknown. Why would a developing system use cell recruitment as a growth mechanism when it can potentially reach any target size by controlling cell proliferation, cell growth, or apoptosis? Recent work in the Drosophila wing shows that cell recruitment is a minor contributor to wing growth, compared to cell proliferation, suggesting that perhaps cell recruitment works a fine-tuning mechanism that provides precision to the final size of the organ [26]. Here, we used a mathematical-modeling approach to drive the hypotheses that the rate of cell proliferation of wing-committed cells is naturally coupled to the speed of recruitment of cells into the wing domain. This dynamic interplay between cell proliferation and recruitment has a direct consequence on the duration of the recruitment process. In particular, when cell proliferation of wing-committed cells is faster or slower than normal, the recruitment process terminates sooner or later than in the wild-type condition, respectively (Fig. 5). This coupling results in a compensatory mechanism that could help the developing wing pouch to reach a robust final size, despite perturbations in cell proliferation rates, a fact that was observed experimentally more than 20 years ago [19], but has not yet been explained mechanistically. Interestingly, this non-linear compensatory mechanism works as a size control mechanism almost as expected analytically using a linear approximation of the model (Fig. 4C,D).

**Figure 5.**
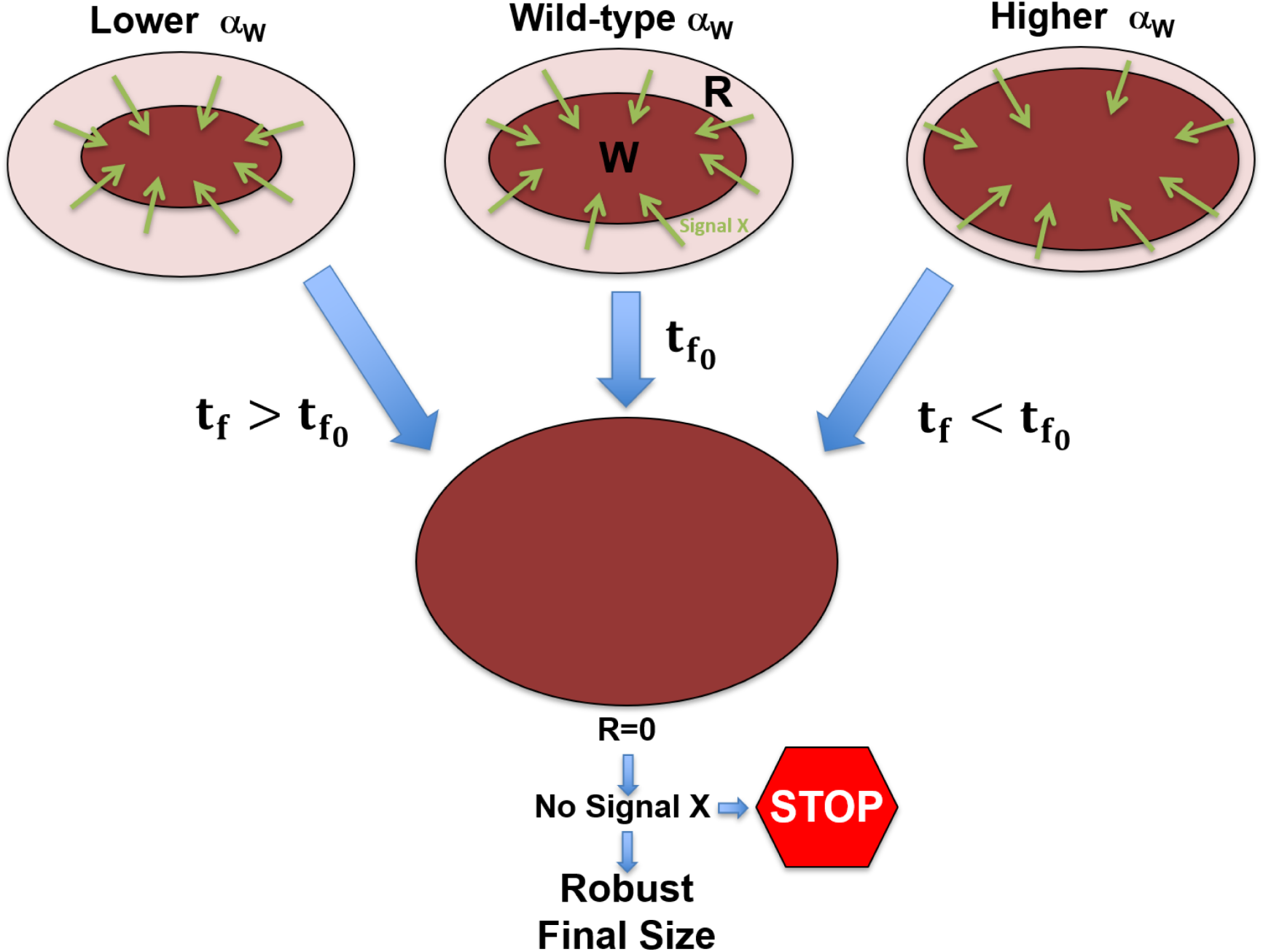
Dynamics of cell proliferation and recruitment control growth of the Drosophila wing disc. W cells (dark red region) increase by cell proliferation and by recruiting neighboring R cells (light red region). Growth is arrested when R cells are completely extinguished, perhaps due to a growth factor signal X that is produced by R cells and sustains cell proliferation in W cells. When the cell proliferation rate of W cells is increased (or decreased), the W domain grows faster (or slower) but R cells are extinguished also faster (or slower) reaching at the end the same final size.

The model relays on some assumptions that deserve further discussion. First, the model assumes a circular geometry, which is essential for growing the number of recruitable cells and accelerate the recruitment process over time. Since W grows exponentially, so does the recruitment boundary (Fig. 5). How much the geometry of the system affects the implications of the model? Importantly, the recruitment function F(W,R) has the same form when the geometry of the system is assumed to be elliptical which resembles more closely the geometry of the wing pouch (see Appendix 4). Second, the model assumes that the disc stops growing when recruitable cells are extinguished. While there is experimental support for this assumption [31], it is left to future studies to provide a definitive proof and eventually a mechanistic explanation of this observation. One possibility is that R cells provide an unidentified chemical growth factor (signal X) that instruct W to proliferate; once R cells are extinguished at the expense of the recruitment process, they also stop producing signal X making the whole system to arrest growth (Fig. 5). Finally, notice that although the mechanism that compensates variations in cell proliferation rates with the duration of growth occurs in a broad range of perturbations (as evidenced by *t_f_* being a decreasing function of α_W_ in Sup. Fig. 3A), it controls well final size, only in a range of perturbations in cell proliferation rates (Sup. Fig. 3B). Thus, it is possible that other mechanisms such as mechanical feedback models [20] or morphogen scaling strategies [33,34] may also be contributing to size control under a broader range of genetic and environmental perturbations.

In summary, our work suggests that a growth control strategy to compensate against fluctuations in cell proliferation rates may be at work in the Drosophila wing disc and provides guidance into how this can be tested experimentally using the genetic toolkit available in Drosophila. Given that cell recruitment is a widespread mechanism in developing organs [32], the mechanism described here provides a more general solution to the problem of how organs robustly attain a reproducible size despite variations in cell proliferation rates, a long-standing problem in developmental biology.

## Acknowledgments

We thank José Luis Fernández-López and Damián Jacinto-Méndez for technical assistance; and Pablo Padilla-Longoria, Yuriria Cortez-Poza, and Eduard De La Cruz-Burelo for discussions and comments on the manuscript.

