## Supplementary Material for "Interplay between cell proliferation and recruitment controls the duration of growth and final size of the Drosophila wing"

**Contents:**

**Appendix 1:** Construction and analysis of recruitment function F(W,R)……………………….. 2

**Supplementary Figure 1.** Geometrical considerations of the mathematical model……..…….. 4

**Appendix 2:** Growth of W is bounded by an exponential function……………..…………….. 5

**Appendix 3:** Analytical behavior of the dynamics of R............................................................. 5

**Supplementary Figure 2.** Error values for the optimization problem………..……..………….. 9

**Supplementary Figure 3.** Temporal growth control is limited to a range of perturbations in $\alpha_W$. 9

**Supplementary Figure 4.** Worst-case scenario in the optimization problem still provides some compensation in $t_f$ against perturbations in $\alpha_W$………...……………………………….……… 10

**Supplementary Figure 5.** Consequences on the dynamics of R when $\alpha_W$ and $\alpha_R$ are perturbed at the same time……………………………..…………………………………………………… 10

**Appendix 4:** Model results are unaffected when the geometry is elliptical…………...…….. 10

**Appendix 1. Construction and analysis of the recruitment function F(W,R)**

Cell recruitment is a process driven by a cell polarization signal driven by cell-to-cell interactions and as such it is a contact and geometry-dependent process (Sup. Fig. 1A). Therefore, the rate of cell recruitment should be proportional to:

i) The number of W cells at the recruitment front, i.e. at the boundary of the W population. Assuming a circular geometry (see Appendix 5 for an elliptical geometry), the area of the W domain is given by:

$$\mathrm{W} \times a = \pi \times \left(\frac{\mathrm{D}}{2}\right)^2 \qquad \text{(A1.1)}$$

where $a$ is the area of an individual W cell (assuming constant for all W cells) and D is the diameter of the W domain. The number of cells at the recruitment front, $\mathrm{W}_p$, is therefore given by the perimeter of the W domain divided by the diameter of an individual W cell, $d$:

$$\mathrm{W}_p = \frac{\pi \mathrm{D}}{d} \qquad \text{(A1.2)}$$

Substituting D from equation (A1.1) into (A1.2) and writing $a = \pi \times \left(\frac{d}{2}\right)^2$ we get (Sup. Fig. 1B):

$$\mathrm{W}_p = \frac{\pi}{d} D = \frac{\pi}{d}\left(d\sqrt{\mathrm{W}}\right) = \pi\sqrt{\mathrm{W}} \qquad \text{(A1.3)}$$

ii) The rate of recruitment depends on available R cells, but this rate is saturated when the number of R cells exceeds the number of recruiter cells at the recruitment front (equation A1.3). To model this saturation process, we use a Hill function:

$$\mathrm{Hill}(\mathrm{W},\mathrm{R}) = \frac{\mathrm{R}}{k\pi\sqrt{\mathrm{W}} + \mathrm{R}} \qquad \text{(A1.4)}$$

Note that when $k$=1, the Hill function is equal to 0.5 when R= $\pi\sqrt{\mathrm{W}}$. However, when $k$<1 we have that the Hill function has a higher level $\left(\frac{1}{1+k}\right)$ when R= $\pi\sqrt{\mathrm{W}}$. We begin the exploration of the dynamics of the system making $k$=1 (Fig. 2), but then we allow it to vary around 1.

iii) The Hill function defined by equation (A1.4) makes the recruitment rate very low as R→0, but we realized that this is unrealistic in practice because there is no reason to think that the recruitment process will slow down dramatically as R decreases. For this reason, we introduced a constant term $\delta$ into the numerator of the Hill function (A1.4):

$$\mathrm{Hill(W,R)} = \frac{\mathrm{R}+\delta}{k\pi\sqrt{\mathrm{W}}+\mathrm{R}} \qquad \text{(A1.5)}$$

Note that if $\delta$ is chosen so that it is always less or equal than $k\pi\sqrt{\mathrm{W}}$ it will only play a significant role when R is small (for this reason we chose $\delta = k\pi\sqrt{\mathrm{W}_0}$ in our simulations). Taken together, we have that the recruitment rate F(W,R) will be given by the product of (A1.3) and (A1.5):

$$\mathrm{F(W,R)} = \rho\pi\sqrt{\mathrm{W}}\,\frac{\mathrm{R}+\delta}{k\pi\sqrt{\mathrm{W}}+\mathrm{R}} \qquad \text{(A1.6)}$$

where $\rho$ is the number of cells that get recruited per unit time for each recruiter cell at the recruitment front.

To make sure that equation (A1.6) behaves as experimentally expected, let us consider the behavior of F(W,R) in the following cases:

Case 1: R$\gg k\pi\sqrt{\mathrm{W}}$

This case applies at the beginning of the early third instar where there are plenty of cells available for recruitment. Notice that the contribution of δ and $\pi\sqrt{\mathrm{W}}$ in the numerator and denominator of the Hill function in (A1.5), respectively, are negligible and $\mathrm{F(W,R)} \approx \rho\pi\sqrt{\mathrm{W}}$. Thus, in this scenario, F(W,R) is simply proportional to the number of recruiter cells.

Case 2: R$\approx \pi\sqrt{\mathrm{W}}$

This case occurs when recruitment dominates over proliferation of R cells until the number of R cells is approximately the number of cells at the recruitment front. Since W is an exponentially increasing function of time $\delta$ becomes negligible in the numerator of the Hill function in (A1.5) and the recruitment rate is $\mathrm{F(W,R)} \approx \frac{1}{k+1}\rho\pi\sqrt{\mathrm{W}}$.

Case 3: R≪ $k\pi\sqrt{W}$

Lastly, when there are not too many cells left for recruitment, R becomes negligible in both the numerator and the denominator of the Hill function in (A1.5). In this case, the recruitment process no longer depends on how many recruiters are available and the recruitment rate approaches a constant rate of $F(W,R) \approx \frac{\rho\delta}{k}$.

**Supplementary Figure 1**

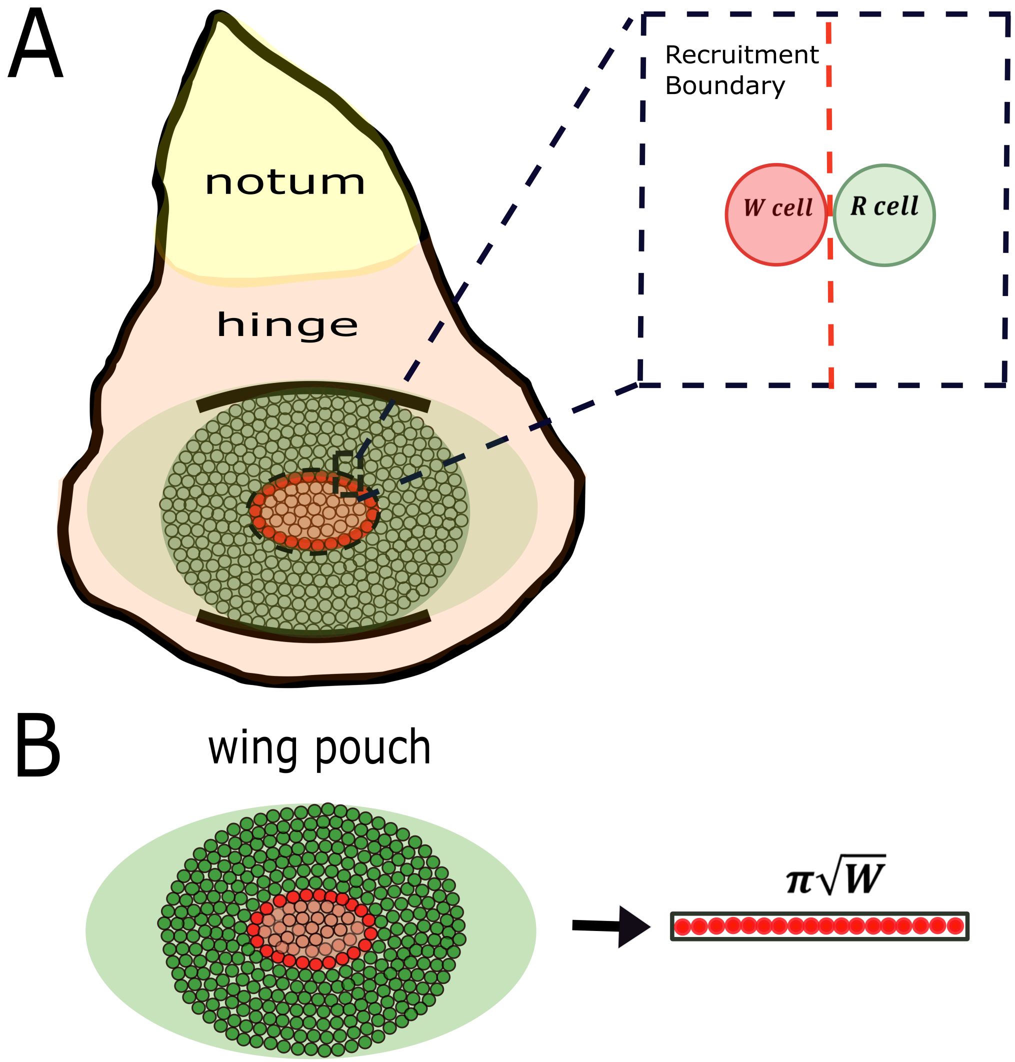


**Supplementary Figure 1**. Geometrical considerations of the mathematical model.
(A) Cartoon depicting a wing imaginal disc. W cells are shown in red and R cells are shown in green. Note that only R cells that are in contact with W cells (bright red) are able to be recruited at that time (inset).
(B) Assuming a circular geometry of the W domain, we can compute the number of recruiter cells at the recruitment boundary $\pi\sqrt{W}$. Note that this number is independent of cell size.

### Appendix 2: Growth of W is bounded by an exponential function

In order to use an exponential approximation of the solution of W in equations (3.1-3.2), we will show that when taken into account cell proliferation + recruitment, growth of W is bounded by an exponential function (Fig. 2A).

Let $\mathrm{U} = \sqrt{\mathrm{W}}$, then $\frac{d\mathrm{U}}{dt} = \frac{1}{2\sqrt{\mathrm{W}}}\frac{d\mathrm{W}}{dt}$. Upon substitution into (3.1) we get:

$$2\mathrm{U}\frac{d\mathrm{U}}{dt} = \alpha_W \mathrm{U}^2 + \rho\pi\mathrm{U}\frac{\mathrm{R}+\delta}{k\pi\mathrm{U}+\mathrm{R}} \qquad \text{(A2.1)}$$

Note that equation (A2.1) only depends on U and R. Dividing by 2U, we get:

$$\frac{d\mathrm{U}}{dt} = \frac{\alpha_W}{2}\mathrm{U} + \frac{\rho\pi}{2}\frac{\mathrm{R}+\delta}{k\pi\mathrm{U}+\mathrm{R}} \qquad \text{(A2.2)}$$

Now, note that since U is a non-decreasing function of time, then $\frac{\mathrm{R}+\delta}{k\pi\mathrm{U}+\mathrm{R}} \leq 1$. Then, an upper bound function of U($t$), $\mathrm{U}_{\mathrm{up}}(t) \geq \mathrm{U}(t)$ satisfies the following linear equation:

$$\frac{d\mathrm{U}}{dt} = \frac{\alpha_{\mathrm{W}}}{2}\mathrm{U} + \frac{\rho\pi}{2} \qquad \text{(A2.3)}$$

Equation (A2.3) can be solved analytically and has the following general solution:

$$\mathrm{U}_{\mathrm{up}}(t) = \mathrm{U}_0 \exp\left(\frac{\alpha_{\mathrm{W}}}{2}t\right) + \beta \qquad \text{(A2.4)}$$

where $\beta = -\frac{\rho\pi}{\alpha_{\mathrm{W}}}$ and $\mathrm{U}_0$ is an integration constant. Then $(\mathrm{U}_{\mathrm{up}}(t))^2 \geq (\mathrm{U}(t))^2 = \mathrm{W}(t)$. Assuming that $(\mathrm{U}_{\mathrm{up}}(0))^2 = \mathrm{W}(0)$, we get that $\mathrm{U}_0 = \sqrt{\mathrm{W}_0} - \beta$. Therefore, an upper bound function of W(t) is given by:

$$\mathrm{W}_{\mathrm{up}}(t) = \left[\left(\sqrt{\mathrm{W}_0} + \frac{\rho\pi}{\alpha_{\mathrm{W}}}\right)\exp\left(\frac{\alpha_{\mathrm{W}}}{2}t\right) - \frac{\rho\pi}{\alpha_{\mathrm{W}}}\right]^2 \qquad \text{(A2.5)}$$

### Appendix 3: Analytical behavior of the dynamics of R

The dynamics of R is expected to depend on parameter values; if cell proliferation dominates over cell recruitment then R is expected to growth indefinitely. Alternatively, if cell recruitment occurs faster than cell proliferation, then R cells will be extinguished at some point. We will prove mathematically (under the assumption that $\alpha = \alpha_{\mathrm{W}} = \alpha_{\mathrm{R}}$) that there are three conditions in parameter space that define the full dynamics of R in the system.

*Case 1*: $k < \frac{\rho}{\alpha}\left(1 + \frac{\delta}{R_0}\right) - \frac{R_0}{\pi\sqrt{W_0}}$

In this case, it is easy to show that $\frac{dR}{dt} < 0$ for all t. Since $\pi\sqrt{W} > 0$, we can multiply the initial inequality by this factor:

$\frac{R_0}{\pi\sqrt{W_0}} + k - \frac{\rho}{\alpha} - \frac{\rho\delta}{\alpha R_0} < 0 \Rightarrow \frac{R_0\sqrt{W}}{\sqrt{W_0}} + \pi\sqrt{W}\left(k - \frac{\rho}{\alpha} - \frac{\rho\delta}{\alpha R_0}\right) < 0$

Since W is increasing, then $\frac{W}{W_0} > 1$. We have that:

$$R_0 + \pi\sqrt{W}\left(k - \frac{\rho}{\alpha} - \frac{\rho\delta}{\alpha R_0}\right) < \frac{R_0\sqrt{W}}{\sqrt{W_0}} + \pi\sqrt{W}\left(k - \frac{\rho}{\alpha} - \frac{\rho\delta}{\alpha R_0}\right) < 0$$

We can then multiply the last inequality by $\frac{\alpha R}{k\pi\sqrt{W}+R}$ (again a positive number) and we get:

$$\alpha R\frac{k\pi\sqrt{W}+R_0}{k\pi\sqrt{W}+R} - \rho\pi\sqrt{W}\frac{R}{k\pi\sqrt{W}+R} - \rho\pi\sqrt{W}\frac{\frac{R}{R_0}\delta}{k\pi\sqrt{W}+R} < 0 \qquad \text{(A3.1)}$$

For $t$=0, the left-hand side of the previous expression is exactly $\frac{dR}{dt}(t = 0)$. Therefore, $\frac{dR}{dt}(t = 0) < 0$.

Consider $t$=$dt$, where $dt$ is a very small non-zero time. Now, since the derivative of R is negative at $t$=0 then, $R(dt) < R_0$ and since the inequality (A3.1) is valid also at $t$=$dt$, we have that:

$$\alpha R(dt)\frac{k\pi\sqrt{W(dt)}+R_0}{k\pi\sqrt{W(dt)}+R(dt)} - \rho\pi\sqrt{W(dt)}\frac{R(dt)}{k\pi\sqrt{W(dt)}+R(dt)} - \rho\pi\sqrt{W(dt)}\frac{\frac{R(dt)}{R_0}\delta}{k\pi\sqrt{W(dt)}+R(dt)} < 0$$

But since $R(dt) < R_0$ then $\frac{k\pi\sqrt{W(dt)}+R_0}{k\pi\sqrt{W(dt)}+R(dt)} > 1$ and $\frac{R(dt)}{R_0} < 1$ hence:

$\frac{dR}{dt}(t = dt) = \alpha R(dt) - \rho\pi\sqrt{W(dt)}\frac{R(dt)}{k\pi\sqrt{W(dt)}+R(dt)} - \rho\pi\sqrt{W(dt)}\frac{\delta}{k\pi\sqrt{W(dt)}+R(dt)} <$

$\alpha R(dt)\frac{k\pi\sqrt{W(dt)}+R_0}{k\pi\sqrt{W(dt)}+R(dt)} - \rho\pi\sqrt{W(dt)}\frac{R(dt)}{k\pi\sqrt{W(dt)}+R(dt)} - \rho\pi\sqrt{W(dt)}\frac{\frac{R(dt)}{R_0}\delta}{k\pi\sqrt{W(dt)}+R(dt)} < 0$

Iterating this process provides an inductive proof that $\frac{dR}{dt} < 0$ for all *t*.

<u>*Case 2*</u>*:* $k > \frac{\rho}{\alpha}\left(1 + \frac{\delta}{R_0}\right)$

In this case, it is easy to show that $\frac{dR}{dt} > 0$ for all *t*:

$k - \frac{\rho}{\alpha}\left(1 + \frac{\delta}{R_0}\right) > 0 \Rightarrow R + \pi\sqrt{W}\left[k - \frac{\rho}{\alpha}\left(1 + \frac{\delta}{R_0}\right)\right] > 0$ (because W and R are positive numbers for all $t<t_f$).

We can then multiply the last inequality by $\frac{\alpha R}{k\pi\sqrt{W}+R}$ (again a positive number) and we get:

$$\alpha R - \rho\pi\sqrt{W}\frac{R\left(1+\frac{\delta}{R_0}\right)}{k\pi\sqrt{W}+R} > 0 \qquad \text{(A3.2)}$$

For *t*=0, the left-hand side of the previous expression is exactly $\frac{dR}{dt}(t = 0)$**.** Therefore, $\frac{dR}{dt}(t = 0) > 0$.

Consider *t*=*dt*, where *dt* is a very small non-zero time. Now, since the derivative of R is positive at *t*=0 then, $R(dt)>R_0$ and since expression (A3.2) is valid also at *t*=*dt*, we have that:

$$\frac{dR}{dt}(t = dt) = R(dt) - \rho\pi\sqrt{W(dt)}\frac{R(dt) + \delta}{k\pi\sqrt{W(dt)} + R(dt)}$$

$$> \alpha R(dt) - \rho\pi\sqrt{W(dt)}\frac{R(dt)\left(1 + \frac{\delta}{R_0}\right)}{k\pi\sqrt{W(dt)} + R(dt)} > 0$$

Iterating this process provides an inductive proof that $\frac{d\mathrm{R}}{dt} > 0$ for all *t*.

<u>*Case 3:*</u> $\frac{\rho}{\alpha}\left(1+\frac{\delta}{\mathrm{R}_0}\right) > k > \frac{\rho}{\alpha}\left(1+\frac{\delta}{\mathrm{R}_0}\right) - \frac{\mathrm{R}_0}{\pi\sqrt{\mathrm{W}_0}}$

In this case, we already showed in Case 1 that $\frac{d\mathrm{R}}{dt}(t=0) > 0$:

$\frac{\mathrm{R}_0}{\pi\sqrt{\mathrm{W}_0}} + k - \frac{\rho}{\alpha} - \frac{\rho\delta}{\alpha\mathrm{R}_0} > 0 \Rightarrow \frac{\mathrm{R}_0\sqrt{\mathrm{W}}}{\sqrt{\mathrm{W}_0}} + \pi\sqrt{\mathrm{W}}\left(k - \frac{\rho}{\alpha} - \frac{\rho\delta}{\alpha\mathrm{R}_0}\right) > 0$ (because W is a positive number for all *t*).

We can then multiply the last inequality by $\frac{\alpha\mathrm{R}}{k\pi\sqrt{\mathrm{W}}+\mathrm{R}}$ (again a positive number) and we get:

$$\alpha\mathrm{R}\frac{k\pi\sqrt{\mathrm{W}}+\frac{\mathrm{R}_0\sqrt{\mathrm{W}}}{\sqrt{\mathrm{W}_0}}}{k\pi\sqrt{\mathrm{W}}+\mathrm{R}} - \rho\pi\sqrt{\mathrm{W}}\frac{\mathrm{R}}{k\pi\sqrt{\mathrm{W}}+\mathrm{R}} - \rho\pi\sqrt{\mathrm{W}}\frac{\frac{\mathrm{R}}{\mathrm{R}_0}\delta}{k\pi\sqrt{\mathrm{W}}+\mathrm{R}} > 0$$

For *t*=0, the left-hand side of the previous expression is exactly $\frac{d\mathrm{R}}{dt}(t=0)$**.** Therefore, $\frac{d\mathrm{R}}{dt}(t=0) > 0$.

In this case, it is not possible to show further what happens to the dynamics of R and it is left to numerical analysis what happens in this scenario (see Fig. 2E). A graphical representation of the findings in this Appendix is shown in Fig. 2D.

### Supplementary Figure 2

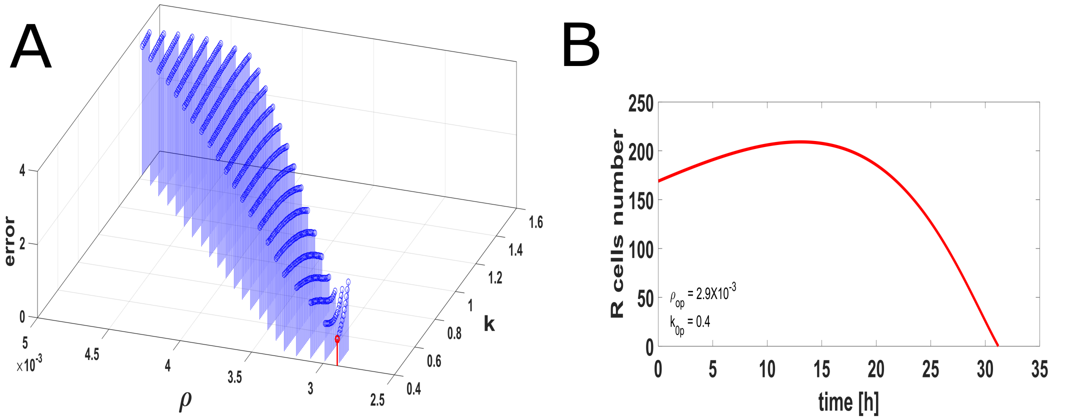


**Supplementary Figure 2**. Error values for the optimization problem.
(A) Optimization error defined as the absolute-value difference between the $t_{f\,ctl}$ and the estimate of $t_f$ (see Fig. 4C) by numerically solving the model equations (3.1-3.2) for different parameter values in a region of parameter space depicted in Fig. 4C (black region) and different perturbations in $\alpha_W$. The red dot is the optimal (minimum error) of the system. Note that the optimal occurs when both $k$ and $\rho$ are small.
(B) Plot of R *vs.* $t$ for $\alpha_{W0}$ and the parameter values corresponding to the optimal in A. Note that for these parameter values, R has a dynamic behavior that is very similar to what is observed experimentally (Fig. 3D, right).

### Supplementary Figure 3

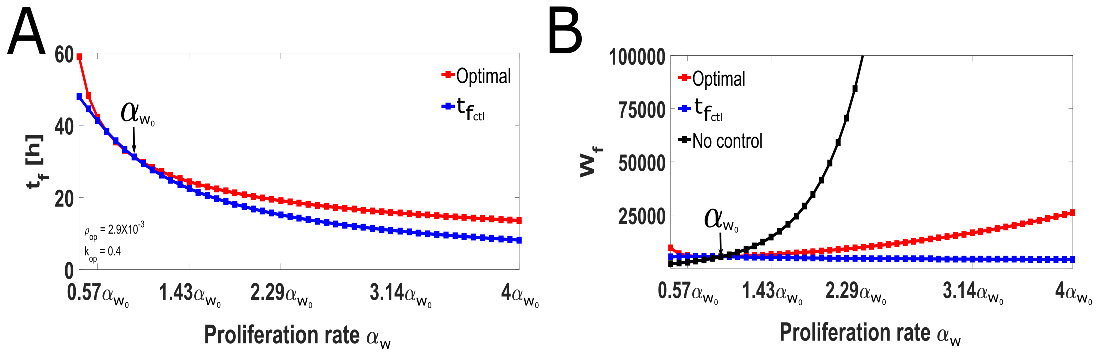


**Supplementary Figure 3**. Temportal growth control is limited to a range of perturbations in $\alpha_W$.
Plots of $t_f$ (A) and $W_f$ (B) *vs.* perturbations in $\alpha_W$ for an extended range of perturbations compared to Fig. 4D and 4E, respectively.

### Supplementary Figure 4

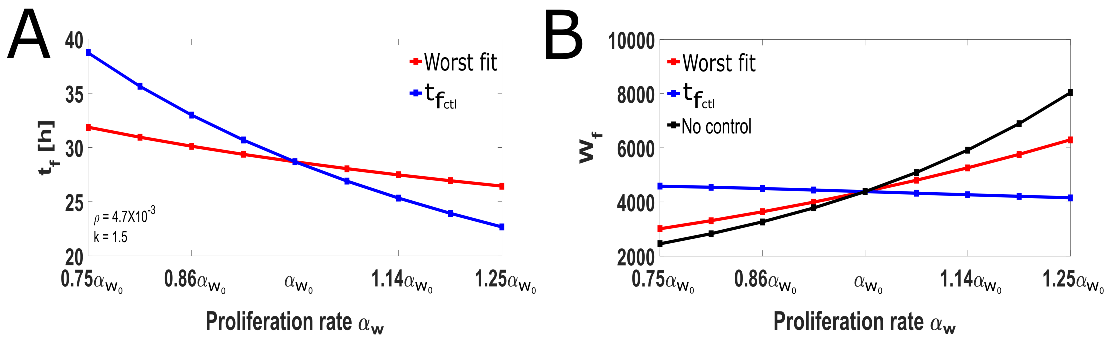


**Supplementary Figure 4**. Worst-case scenario in the optimization problem still provides some compensation in $t_f$ against perturbations in $\alpha_W$.
Plots of $t_f$ (A) and $W_f$ (B) *vs.* perturbations in $\alpha_W$ for the parameter values that maximize the error in Sup Fig. 2A. Note that $t_f$ is still a decreasing function of $\alpha_W$ and $W_f$ is more controlled with respect to the curve without any control (compare red and black curves in B).

### Supplementary Figure 5

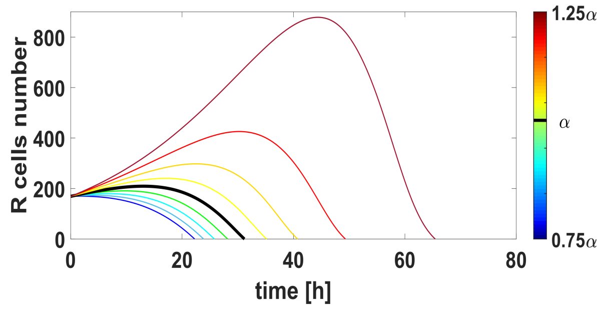


**Supplementary Figure 5**. Consequences on the dynamics of R when $\alpha_W$ and $\alpha_R$ are perturbed at the same time.
Temporal dynamics of the R population when $\alpha_W$ and $\alpha_R$ are perturbed simultaneously. Note that in this case the time in which R=0 is an increasing function of $\alpha$. Colors correspond to the interval of perturbations in $\alpha$. The simulations were obtained using the optimal parameters showed in Fig. 4D.

#### Appendix 5: Model results are unaffected when the geometry is elliptical

Our model assumes that the wing pouch is a circular disc and the recruitment front as a circle. However, in reality the geometry of the system is approximately elliptical. Here we show that the model is effectively unchanged when the recruitment front is assumed to be elliptical.

Let us suppose that the area of the wing blade is an ellipse with semi-mayor axis $a$ and semi-minor axis $b$, so the area and the perimeter of the ellipse are given by:

$$A = \pi ab \ , \ P \approx 2\pi\sqrt{\frac{a^2+b^2}{2}} \qquad \text{(A4.1)}$$

Now, let us consider that the W and R cells can have many contacts each other; then, we can suppose that the cells are identical polygons of side $n$ inscribed in a circle of radius $r$ and area:

$$A_p = \frac{1}{2}nr^2 \sin\left(\frac{2\pi}{n}\right)$$

Thus, we can define the ellipse area in terms of each cell as follows:

$$A = \mathrm{W}A_p = \frac{1}{2}nr^2 \sin\left(\frac{2\pi}{n}\right) \qquad \text{(A4.2)}$$

Matching the equations (A4.1) and (A4.2), we can express the radius of each cell as a function of the total number of W cells:

$$r = \sqrt{\frac{2\pi ab}{W(t)n sin(\frac{2\pi}{n})}}$$

To take on account the number of W cells that can recruit, we need to estimate the number of W cells that are in the recruitment front $\mathrm{W}_p$, this quantity is equal to the perimeter of the ellipse divided by the diameter of each cell:

$$\mathrm{W}_p = \frac{P}{2r} \approx \frac{2\pi\sqrt{\frac{a^2+b^2}{2}}}{2\sqrt{\frac{2\pi ab}{\mathrm{W}n sin(\frac{2\pi}{n})}}}$$

$$\Rightarrow \qquad \mathrm{W}_p \approx \pi\sqrt{\mathrm{W}}\sqrt{\frac{(a^2+b^2)n sin\left(\frac{2\pi}{n}\right)}{4\pi ab}} \qquad \text{(A4.3)}$$

Therefore, the dynamic of the model assuming that the recruitment front is an ellipse is governed only by the behavior of $\pi\sqrt{\mathrm{W}}$.
